# Viewing communities as coupled oscillators: elementary forms from Lotka and Volterra to Kuramoto

**DOI:** 10.1101/2020.05.26.112227

**Authors:** Zachary Hajian-Forooshani, John Vandermeer

## Abstract

Ecosystems and their embedded ecological communities are almost always by definition collections of oscillating populations. This is apparent given the qualitative reality that oscillations emerge from consumer-resource interactions, which are the simple building blocks for ecological communities. It is also likely always the case that oscillatory consumer-resource pairs will be connected to one another via trophic cross-feeding with shared resources or via competitive interactions among resources. Thus, one approach to understanding the dynamics of communities conceptualizes them as collections of oscillators coupled in various arrangements. Here we look to the pioneering work of Kuramoto on coupled oscillators and ask to what extent can his insights and approaches be translated to ecological systems. We explore all possible coupling arrangements of the simple case of three oscillator systems with both the Kuramoto model and with the classical Lotka-Volterra equations that are foundational to ecology. Our results show that the six-dimensional analogous Lotka-Volterra systems behave strikingly similarly to that of the corresponding Kuramoto systems across all possible coupling combinations. This qualitative similarity in the results between these two approaches suggests that a vast literature on coupled oscillators that has largely been ignored by ecologists may in fact be relevant in furthering our understanding of ecosystem and community organization.

## Introduction

Within ecological communities, it is undeniable that interacting species assemblages are composed of pairs of consumers and their resources which, by definition makes them collections of oscillators. To the extent that the consumers tend to overlap in their diets, or the resources interact with one another, these assemblages may be thought of as systems of coupled oscillators. These consumer-resource oscillators necessarily act as fundamental building blocks which scale up to communities and ecosystems. Although ecologists have long been interested in understanding large assemblages of interacting species, relatively little research in ecology has drawn on the body of theory associated with coupled oscillators. In many branches of science the issue of coupled oscillators is a key metaphor for developing general theory, from electronics to neurobiology (Norton et al., 2018; Laing, 2017; Fukuyama, and Okugawa, 2017).

One elegant perspective on coupled oscillators is the abstraction of Kuramoto (1975; 1984) which, in the present context, takes each resource consumer pair and assumes it forms a limit cycle and that connections among the limit cycles is such that they will tend to phase synchronize. Kuramoto’s classical model is formulated on the circle and the oscillator conditions indicated as the angle Θ made by the point of resource and consumer on the unit circle, taken to represent the limit cycle of the oscillator. Presuming that synchronization will occur, Kuramoto writes:

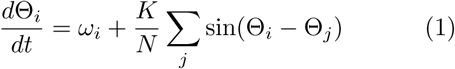

where *ω*_*i*_ is the winding number of oscillator *i* (the rate of advancement on the circle dictated by the inherent oscillations), *K* is the intensity of coupling, and *N* is the number of oscillators. Clearly the intent of the model is to view the phase of the oscillations (not the amplitude) as the key dynamical force. All oscillators are identical (with the possible exception of the winding number) and couplings are taken to be universal (all to all). A rather remarkable result emerges from this simple model – with random initiations, no synchrony occurs when coupling intensities are small, but a critical point of coupling intensity is reached where rapid attainment of synchrony of all oscillators is achieved. This model has been useful in studying large systems of coupled oscillators (Rodrigues, et al., 2016).

Based on the reality of coupling types in ecological consumer-resource systems, there are two general ways in which consumer-resource pairs can be coupled. First, when two consumers share two resources they are, by definition coupled with one another via trophic cross-feeding. If their cross-feeding is relatively weak, they will converge on a pattern of relative in-phase synchrony with one another (Vandermeer, 2004). On the other hand, if coupling is through the competitive interactions of the resources, the oscillators will converge on a pattern of relative anti-phase synchrony with one another (Vandermeer, 2004). Here we refer to these two qualitatively different forms of consumer-resource pair couplings as trophic-coupling and resource-coupling respectively (Figure 1.). The inevitable oscillatory dynamics that emerge from these qualitative coupling arrangements have been confirmed empirically in at least one case (Benincà et al., 2009).

**Figure 1:**
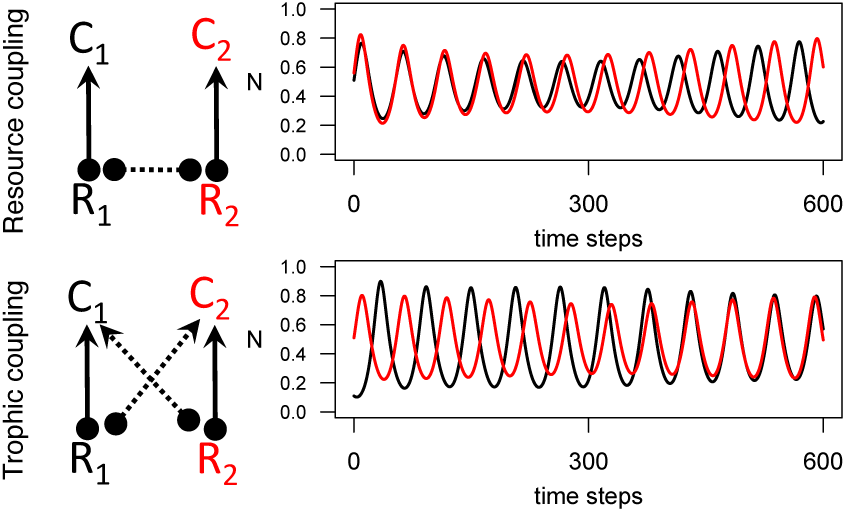
Shows the two qualitative coupling arrangements for consumer-resource oscillators and their dynamic outcomes. As Vandermeer (2004) pointed out, resource coupling (competition between resources) leads to asynchrony and trophic coupling (cross-feeding) leads to synchrony. Note that circles represent negative effects and arrows positive effects and dotted lines emphasize oscillator coupling.

Although it is apparent to ecologists that oscillations are an essential feature that results from the most elementary of ecological interactions, approaches used in the field of complex systems, like those pioneered by Kuramoto, have gained relatively little traction in the field of ecology. It is most frequently the practice in ecology, especially in the foodweb literature, to couple together large networks of ordinary differential equations (e.g. Lotka-Volterra)representing individual populations of consumers and resources. Although this approach has been fruitful, it sometimes leads to unwieldy parameterization, limiting analytical questions to those amenable to linear stability analyses. To explore the potential usefulness of employing approaches such as those of Kuramoto, we here study the concordance between his model and the classical Lotka-Volterra models used in ecology, for the most elementary formulation of an ecological community.

## Methods

Modifying Kuramoto’s model, we write:

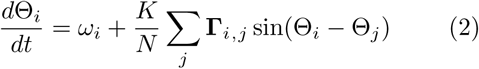

where Kuramoto’s mean field approach has been disaggregated with the adjacency matrix **Γ** stipulating the coupling of each pair of oscillators. Note that **Γ**_*i,j*_ > 0 indicates the oscillators *i* and *j* will synchronize “in phase” while **Γ**_*i,j*_ < 0 indicates they will synchronize “anti-phase.” If we stipulate that **|Γ**_*i,j*|_ = 1.0, the sum of the upper triangle of the adjacency matrix 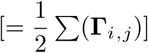 can be −3, −1, 1, or 3 for a three oscillator system. Figure 2 illustrates the basic combinations of a three oscillator system with expected outcomes of oscillator phases based on coupling.

Taking the classic Lotka-Volterra consumer resource equations we write:

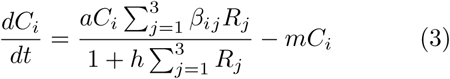

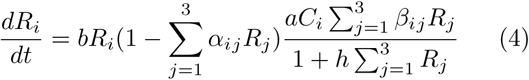

**Figure 2:**
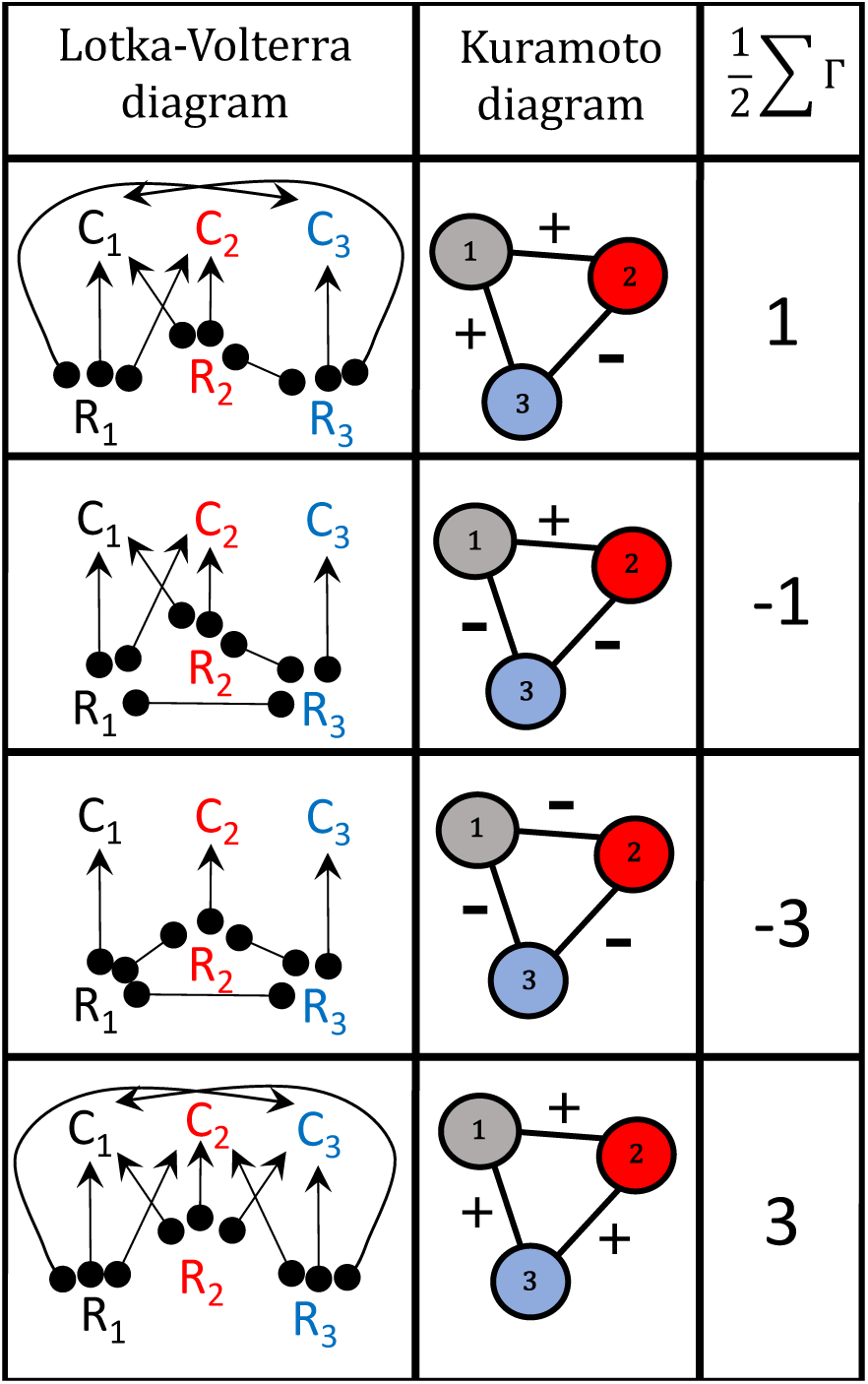
Diagrammatic representation of the analogous forms of Lotka-Volterra and Kuamoto for three oscillator communities. **Lotka-Volterra diagram**: Illustrates the core idea of three consumer/resource coupled oscillators, *C*_*i*_ is the biomass of the *ith* consumer and *R*_*i*_ is the biomass of the *ith* resource. Connectors indicate a positive effect with an arrowhead and a negative effect with a closed circle. **Kuramoto diagram**: illustrates the three oscillators as nodes in a graph and their connections, edges, with the elements of the adjacency matrix (+=1 or = −1) indicated near the edges. 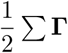 shows the sum of the elements of half of the adjacency matrix to simply represent the four unique coupling arrangements. Note that the coloring scheme of the oscillators is consistent throughout the article.

Where *C* and *R* denote “consumers” and “resources” respectively and *i* ranges from 1 to 3. The basic parameters of the model are: *a* = the attack rate of the consumer, *m* = the mortality rate of the consumer, *h* = the functional response term of the consumer, and *b* = the birth rate of the resource. The parameter *α* represents the strength of competition (resource coupling) between resources and *β* represents the strength of cross-feeding (trophic coupling). The long form of the equations along with the initial conditions for simluations can be found in the Appendix.

In the spirit of Kuramoto’s model, we first located parameter space where individual consumer-resource pairs oscillate in limit cycles (equations 3 and 4). For all simulations presented here, those parameters are: *a* = 0.7, *m* = 0.1, *h* = 3.0 and *b* = 0.3. Given a persistent oscillator in the Lotka-Volterra formulation, we then couple them in the four possible combinations outlined in Figure 2 in the spirit of Vandermeer (2004), where trophic-coupling implies eventual synchrony and resource-coupling implies asynchrony. Manipulating *α*_*ij*_ and *β*_*ij*_ in equations (3) and (4), we create the parameter states for all four coupling arrangements depicted in Figure 2. For all simulations presented here we used low values of coupling coefficients (*β* = 0.01 and *α* = 0.1).

## Results

Employing the Kuramoto model (equation 2) if Σ_*j*_ **Γ**_*i,j*_ = 3, the system synchronizes in phase (Figure 3), if it is −3 the system synchronizes anti-phase, if it is −1 two oscillators are in phase and the third is anti-phase with the two in-phase oscillators. However, if the sum is 1 an intransitive situation emerges in which oscillator 3 is in synchrony with oscillator 1, oscillator 1 is in synchrony with oscillator 2, while oscillator 2 is anti-synchronous with oscillator 3, but, qualitatively stable with all three oscillators wandering around the state space according to the winding number, but separated from one another by 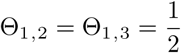 and 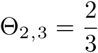. The four qualitativeresults in circular phase space are presented in Figure 3.

**Figure 3:**
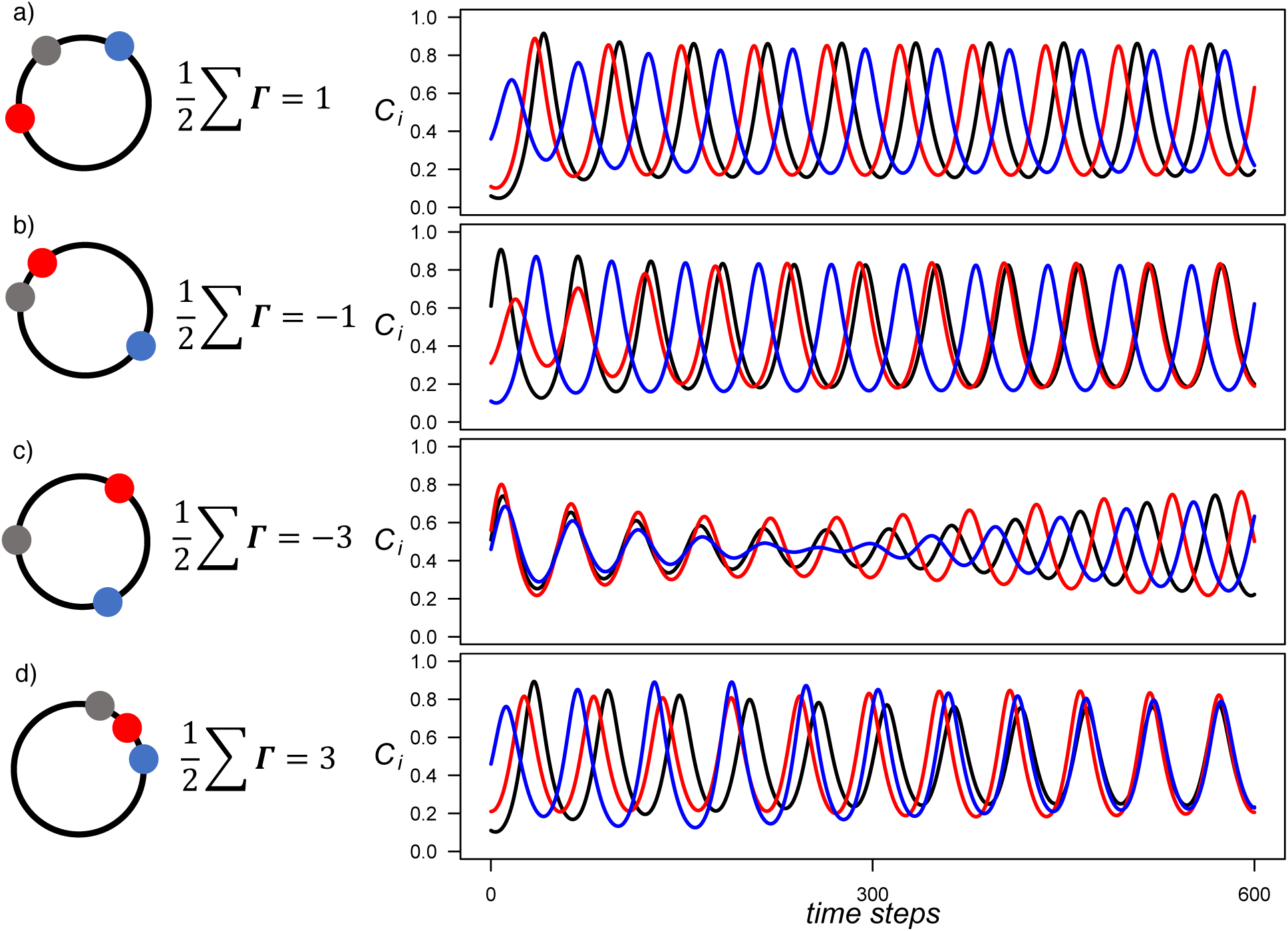
Time series from the four separate coupling configurations of the three dimensional Kuramoto and six dimensional Lotka-Volterra systems. a)-d) show the oscillator phases from the Kuramoto systems and the time series for the analogous six dimensional Lotka-Volterra systems. The clear concordance between the Lotka-Volterra time series and the Kuramoto oscillator phases is evident by the end of the Lotka-Volterra time series plots. Note that only the consumers are plotted in the time series to more clearly demonstrate the correspondence in oscillatory dynamics between LV and Kuramoto. The parameters for all Lotka-Volterra simulations are: *a* = 0.7, *m* = 0.1, *h* = 3.0, *b* = 0.3, *β* = 0.01 and *α* = 0.1. For the Kuramoto model simulations *k* = 0.1.

Employing the LV model (equations 3 & 4), typical time series results of all four ODE simulations are presented in Figure 3. It is clear that the L-V predictions for all four qualitatively distinct cases (Figure 2) are precisely what we get from the simpler Kuramoto approach.

## Discussion

The ubiquity of oscillatory dynamics in ecology has long been appreciated (Blasius et al., 2020). Empirically, across a range of spatiotemporal scales from large scale dynamics of the hare-lynx system (Blasius et al., 1999) to the microcosm experiments of Huffaker (1958), and theoretically emerging from the simplest conceptualizations of consumer-resource interactions (Lotka, 1928; Volterra, 1926), synchronization of coupled consumer resource oscillators is well-known (Vandermeer, 2006). Here we demonstrate that for the four most obvious qualitatively distinct yet ecologically significant coupling patterns in a six-species community (three oscillators), weak coupling leads to precisely the pattern predicted by Kuramoto’s phase coupled system. These results suggest a wholly distinct vision of ecological communities in which the “agents” are not population densities, but rather oscillators.

By reorienting the focus of ecological analogy from individual populations to collections of oscillators, the dynamical nature of the system becomes the central focus rather than questions of stability or persistence. As in other scientific fields, the collective dynamics of these coupled oscillators can provide a useful heuristic for exploring the general properties of large and complex systems that ecologists have long cited with awe (Lawton, 1999; Vellend, 2010). Furthermore, by highlighting the ability to move between classical models in ecology and classical models in the coupled oscillator literature, we suggest that both approaches can be used in tandem and exploited for their strengths. Approaches á la Kuramoto effectively increase the tractability of large complex systems by halving the dimensionality and providing an elegant and intuitive way to visualize the oscillatory dynamics, while approaches á la Lotka-Volterra permit investigation of how basic biological parameters influence dynamics. The most obvious utility of such an approach is where synchronous dynamics are the focus of investigation (e.g. Earn et al.,1998; Blasius et al., 199; Liebhold et al., 2004), and may have practical implications for the management of fisheries (Kaemingk et al., 2018), the planning complex biological control systems in agroecosystems (Vandermeer et al., 2019), or conservation (Earn et al., 2000).

The once popular idea that ecosystems are at, or moving towards, Lyapunov stability is considered passé (e.g., Morozov et al., 2019). The growing appreciation amongst ecologists that ecosystems and communities are dominated by nonlinear processes often outside of equilibrium (Rominger, et al., 2017) suggests that our tool kits to understand ecosystems need to evolve along with our analogies of them. We suggest that networks of oscillators, rather than networks of populations, represent a potentially new paradigm for the examination of ecological communities.

## Acknowledgements

The students in the “Complex Systems in Ecology” course of Fall 2019 at University of Michigan participated in stimulating discussions related to the ideas presented here. Chatura Vaidya and Kristel Sanchez provided feedback on an earlier draft of the manuscript. Several examples from Ferdinand LaMothe and Oscar Peterson gave some intuition into the dynamics of coupled oscillators that were useful in the development of this manuscript. This work supported by NSF grant number DEB – 1853261

## Appendix

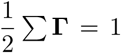 for the Lotka-Volterra system of ordinary differential equations has *C*_1_ with trophic-coupling on *R*_2_ and *R*_3_, *C*_2_ with trophic-coupling on *R*_1_, *C*_3_ with trophic-coupling on *R*_1_, and *R*_2_ and *R*_3_ resource coupled. Initial conditions for Figure 2 are: *R*_1_ = 0.05, *R*_2_ = 0.10, *R*_3_ = 0.35, *C*_1_ = 0.06, *C*_2_ = 0.11, *C*_3_ = 0.36

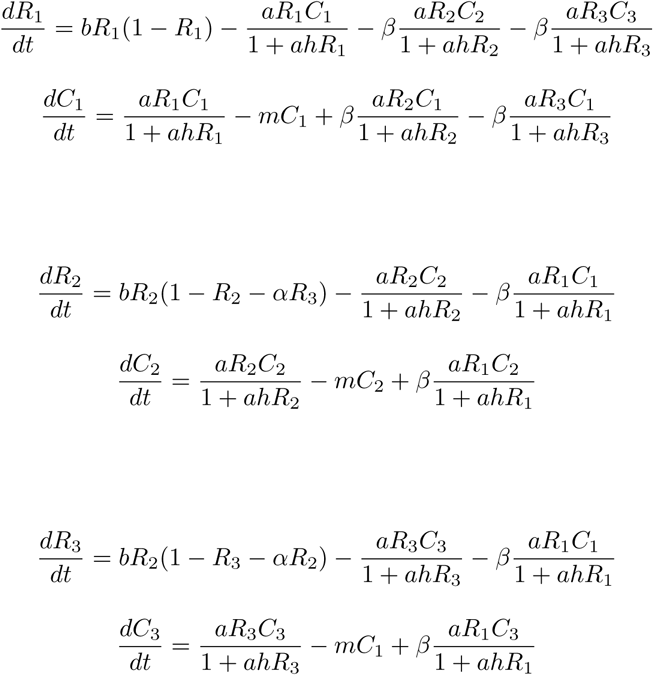

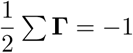 for the Lotka-Volterra system of ordinary differential equations has *C* with trophic-coupling on *R*_2_; *C*_2_ with trophic-coupling on *R*_1_; *R*_2_ and *R*_3_ with resource-coupling and *R*_1_ and *R*_3_ with resourcecoupling. Initial conditions for Figure 2 are: *R*_1_ = 0.60, *R*_2_ = 0.30, *R*_3_ = 0.10, *C*_1_ = 0.61, *C*_2_ = 0.31, *C*_3_ = 0.11

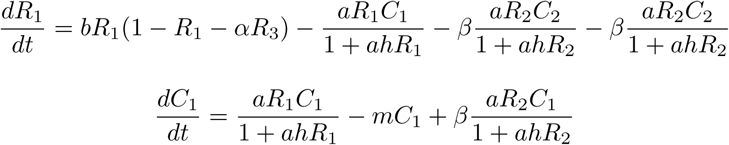

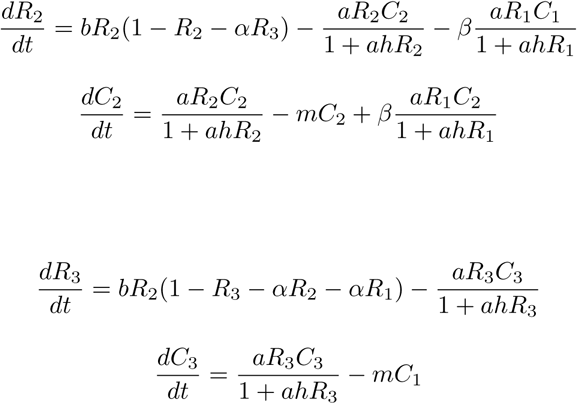

for 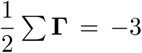 has pairwise resource-coupling between all resources and no trophic-coupling. Initial conditions for Figure 2 are:*R*_1_ = 0.50, *R*_2_ = 0.55, *R*_3_ = 0.45, *C*_1_ = 0.51, *C*_2_ = 0.56, *C*_3_ = 0.46

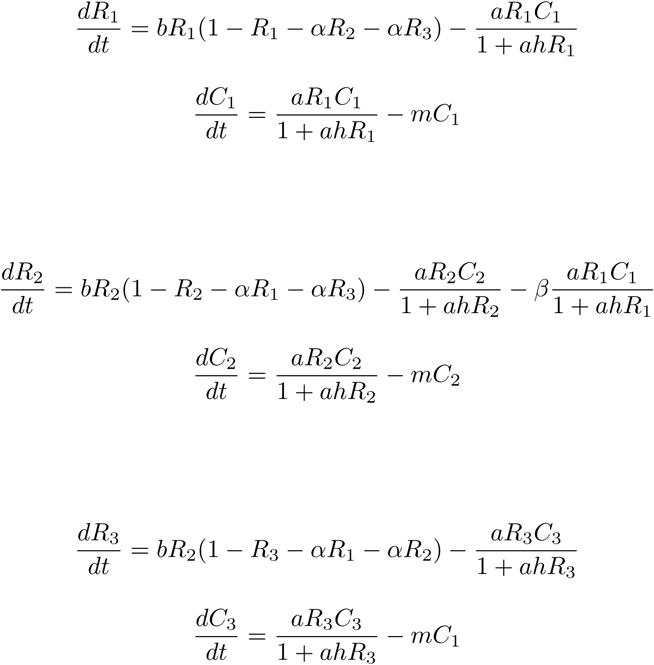

for 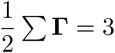 has pairwise trophic-coupling between all consumers and resources and no resource-coupling. Initial conditions for Figure 2 are:*R*_1_ = 0.10, *R*_2_ = 0.25, *R*_3_ = 0.45, *C*_1_ = 0.11, *C*_2_ = 0.21, *C*_3_ = 0.46

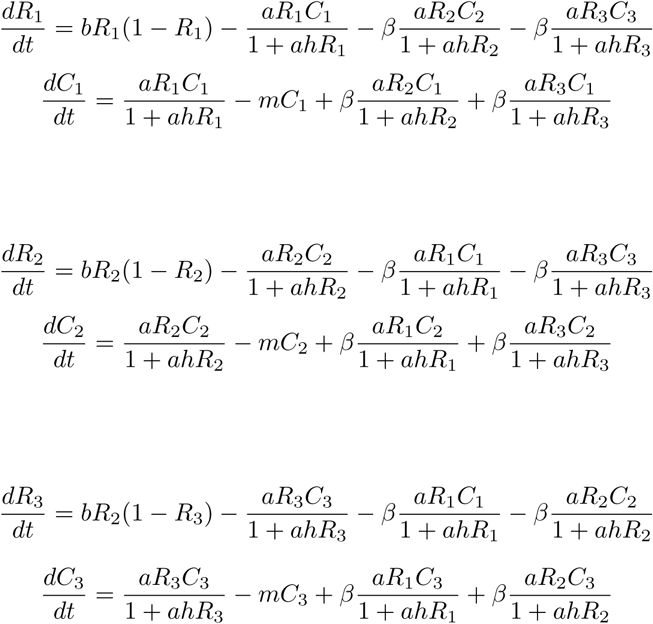

## References

1. Benincà, E., Jöhnk, K. D., Heerkloss, R., Huisman, J. (2009). Coupled predator–prey oscillations in a chaotic food web. Ecology letters, 12(12), 1367–1378.

2. Blasius, B., Huppert, A. and Stone, L., 1999. Complex dynamics and phase synchronization in spatially extended ecological systems. Nature, 399(6734), pp.354–359.

3. Blasius, B., Rudolf, L., Weithoff, G., Gaedke, U. and Fussmann, G.F., 2020. Long-term cyclic persistence in an experimental predator–prey system. Nature, 577(7789), pp.226–230.

4. Earn, D. J., Levin, S. A., & Rohani, P. (2000). Coherence and conservation. Science, 290(5495), 1360–1364.

5. Earn, D. J., Rohani, P., & Grenfell, B. T. (1998). Persistence, chaos and synchrony in ecology and epidemiology. Proceedings of the Royal Society of London. Series B: Biological Sciences, 265(1390), 7–10.

6. Elton, C., & Nicholson, M. (1942). The ten-year cycle in numbers of the lynx in Canada. The Journal of Animal Ecology, 215–244.

7. Fukuyama, T. and Okugawa, M., 2017. Dynamic characterization of coupled nonlinear oscillators caused by the instability of ionization waves. Physics of Plasmas, 24(3), p.032302.

8. Kaemingk, M. A., Chizinski, C. J., Hurley, K. L., & Pope, K. L. (2018). Synchrony—An emergent property of recreational fisheries. Journal of Applied Ecology, 55(6), 2986–2996.

9. Kuramoto, Y. (1975). Self-entrainment of a population of coupled non-linear oscillators. In International symposium on mathematical problems in theoretical physics (pp. 420–422). Springer, Berlin, Heidelberg.

10. Kuramoto, Y. (1984). Cooperative dynamics of oscillator communitya study based on lattice of rings. Progress of Theoretical Physics Supplement, 79, 223–240.

11. Laing, C.R., 2017. Phase oscillator network models of brain dynamics. Computational models of brain and behavior, pp.505–517.

12. Lawton, J.H., 1999. Are there general laws in ecology?. Oikos, pp.177–192.

13. Liebhold, A., Koenig, W. D., & Bjørnstad, O. N. (2004). Spatial synchrony in population dynamics. Annu. Rev. Ecol. Evol. Syst., 35, 467–490.

14. Morozov, A., Abbott, K., Cuddington, K., Francis, T., Gellner, G., Hastings, A., Lai, Y.C., Petrovskii, S., Scranton, K. and Zeeman, M.L., 2019. Long transients in ecology: theory and applications. Physics of Life Reviews.

15. Norton, M., Hunter, I., Moustaka, M., Crisholm, A., Hagan, M., Fahmy, Y. and Fraden, S., 2018. Multistable Dynamical Network of Diffusively Coupled Chemical Oscillators. Bulletin of the American Physical Society, 63.

16. Rodrigues, F.A., Peron, T.K.D., Ji, P. and Kurths, J., 2016. The Kuramoto model in complex networks. Physics Reports, 610, pp.1–98.

17. Rominger, A.J., Overcast, I., Krehenwinkel, H., Gillespie, R.G., Harte, J. and Hickerson, M.J., 2017. Linking evolutionary and ecological theory illuminates non-equilibrium biodiversity. arXiv preprint arXiv:1705.04725.

18. Vandermeer, J. (1993). Loose coupling of predator-prey cycles: entrainment, chaos, and intermittency in the classic MacArthur consumer-resource equations. The American Naturalist, 141(5), 687–716.

19. Vandermeer, J. (2004). Coupled oscillations in food webs: balancing competition and mutualism in simple ecological models. The American Naturalist, 163(6), 857–867.

20. Vandermeer, J. (2006). Oscillating populations and biodiversity maintenance. Bioscience, 56(12), 967–975.

21. Vandermeer, J., Armbrecht, I., de la Mora, A., Ennis, K.K., Fitch, G., Gnthier, D. J., Hajian-Forooshani, Z., Hsun-Yi, H., Iverson, A., Jackson, D., Jha, S., Jiménez-Soto, E., Lopez-Bautista, G., Larsen, A., Li, K., Liere, H., MacDonald, A., Marin, L., Mathis, K. A., Monagan, I., Morris, J. R., Ong, T., Pardee, G. L., Saraeny Rivera-Salinas, I., Vaiyda, C., Williams-Guillen, K., Yitbarek, S., Uno, S., Zeminick, A., Philpott, S. M., Perfecto, I., 2019. The community ecology of herbivore regulation in an agroe-cosystem: Lessons from Complex Systems. BioScience, 69(12), pp.974–996.

22. Vellend, M., 2010. Conceptual synthesis in community ecology. The Quarterly review of biology, 85(2), pp.183–206.

